# Responses of chlorophyll fluorescence to CO_2_ elimination as an indicator of Crassulacean acid metabolism photosynthesis

**DOI:** 10.1101/2024.05.25.595861

**Authors:** Sae Bekki, Kenji Suetsugu, Koichi Kobayashi

**Affiliations:** Department of Biology, Graduate School of Science, Osaka Metropolitan University, 1-1 Gakuen-cho, Naka-ku,Sakai, Osaka 599-8531, Japan; Department of Biology, Graduate School of Science, Kobe University, 1-1 Rokkodai, Nada-ku, Kobe, Hyogo 657-8501, Japan; The Institute for Advanced Research, Kobe University, 1-1Rokkodai, Nada-ku, Kobe, Hyogo 657-8501, Japan; Faculty of Liberal Arts, Science and Global Education, Osaka Metropolitan University, 1-1 Gakuen-cho, Naka-ku, Sakai, Osaka 599-8531, Japan

**Keywords:** *Bulbophyllum drymoglossum*, *Bulbophyllum inconspicuum*, chlorophyll fluorescence, Crassulacean acid metabolism, epiphytic orchids, *Gastrochilus japonicus*, *Kalanchoe pinnata*, *Kalanchoe daigremontiana*, malate, *Oberonia japonica*

## Abstract

Crassulacean acid metabolism (CAM) is found in a wide variety of vascular plant species, mainly those inhabiting water-limited environments. Identifying and characterizing diverse CAM species enhances our understanding of the physiological, ecological, and evolutionary significance of CAM photosynthesis. In this study, we examined the effect of CO_2_ elimination on chlorophyll fluorescence-based photosynthetic parameters in two constitutive CAM *Kalanchoe* species and six orchids. In CAM-performing *Kalanchoe* species, the effective quantum yield of photosystem II showed no change in response to CO_2_ elimination during the daytime but decreased with CO_2_ elimination at dusk. We applied this method to reveal the photosynthetic mode of epiphytic orchids and found that *Gastrochilus japonicus*, *Oberonia japonica*, and *Bulbophyllum inconspicuum*, but not *Bulbophyllum drymoglossum*, are constitutive CAM. Although *B. drymoglossum* had relatively high malate content in leaves, they did not depend on it to perform photosynthesis even under water deficient or high light conditions. Anatomical comparisons revealed a notable difference in the leaf structure between *B. drymoglossum* and *B. inconspicuum*; *B. drymoglossum* leaves possess the large water storage tissue internally, unlike *B. inconspicuum* leaves, which develop pseudobulbs. Our data propose a novel approach to identify and characterize CAM plants without labor-intensive experimental procedures.

**Highlight:** Responses of chlorophyll fluorescence-based photosynthetic parameters to CO_2_ elimination differ between Crassulacean acid metabolism (CAM) and C_3_ metabolism, proposing a novel approach to identify and characterize CAM plants.

## Introduction

Crassulacean acid metabolism (CAM) is a specialized mode of photosynthetic carbon assimilation, in which C_4_ carboxylation via phosphoenolpyruvate carboxylase (PEPC) and C_3_ carboxylation via Rubisco are temporally separated between night and day. The typical CAM cycle can be divided into four phases (Osmond, 1978). At night, CO_2_ is taken into leaves through the stomata, fixed via PEPC, and stored in the vacuole mainly as malate (phase I). At the beginning of the light phase, atmospheric CO_2_ via open stomata is fixed by Rubisco in addition to PEPC (phase II). With stomata being closed in the middle of the day, CO_2_ is released from the decarboxylation of malate and refixed by Rubisco (phase III). At the end of the daytime, malate content decreases, and stomata re-open to take up atmospheric CO_2_, which is fixed mainly by Rubisco (phase IV).

The CAM cycle, including CO_2_ fixation at night and stomatal closure during the daytime, increases water-use efficiency and tolerance to drought stress (Hultine *et al*., 2019). CAM is often observed in vascular plants that inhabit water-limited environments, including desert succulents and tropical epiphytes. Constitutive CAM plants continuously operate CAM photosynthesis in mature tissues, although many of them perform C_3_ photosynthesis when the tissues are young (Winter, 2019). Other CAM species, called facultative CAM plants, reversibly change the mode of photosynthesis between CAM and C_3_ or C_4_ in response to environmental stresses, typically drought stress. In addition, some aquatic species are known to operate CAM photosynthesis to cope with ambient limitations of carbon in aquatic environments (Keeley, 1998). As a result of multiple independent evolution in pteridophytes, gymnosperms, and angiosperms, CAM is estimated to be expressed in ∼6% of vascular plant species (Smith and Winter, 1996).

Identification and characterization of diverse CAM species provide deeper insights into the evolution and significance of CAM photosynthesis. Several approaches have been conducted to identify and investigate CAM plants. Nocturnal acidification is a typical feature specific to CAM species, and the measurement of titratable acidity of soluble extracts from plant tissues is commonly used to identify CAM species (Winter and Smith, 2022). Alternatively, the measurement of malate content at dawn and dusk directly suggests the activity of the CAM cycle. Determination of CO_2_ assimilation by gas exchange measurement or ^14^CO_2_ labeling indicates a relationship between CO_2_ uptake and the light-dark cycle, with nocturnal CO_2_ fixation being a typical feature of CAM photosynthesis (Nishida, 1978; Winter, 2019; Dodd *et al*., 2002). In field-scale assessments, the ^13^C- to-^12^C ratio (δ^13^C), which is associated with the different discrimination against ^13^CO_2_ by Rubisco and PEPC, is used as a signature of CAM plants (Cernusak *et al*., 2013; Messerschmid *et al*., 2021). Other approaches include determination of stomatal openness (Cockburn *et al*., 1979; Males and Griffiths, 2017), activity of CAM-associated enzymes (Dittrich *et al*., 1973; Nimmo, 2000; Dittrich, 1976; Ceusters *et al*., 2021), anatomical investigation (Herrera, 2020; Gilman *et al*., 2024; Barrera Zambrano *et al*., 2014; Heyduk *et al*., 2016), and several omics analyses (Yang *et al*., 2017; Cushman *et al*., 2008; Li *et al*., 2023; Heyduk *et al*., 2019; Abraham *et al*., 2020).

We recently reported that the epiphytic orchid *Thrixspermum japonicum* performs CAM photosynthesis in the leaves but not the roots (Suetsugu *et al*., 2023). Similar to *T. japonicum* leaves, the shootless epiphytic orchid *Taeniophyllum aphyllum* showed nocturnal ^14^CO_2_ uptake and malate accumulation in its roots, suggesting that *T. aphyllum* roots perform CAM photosynthesis as do *T. japonicum* leaves. To confirm this result, we conducted an imaging analysis of pulse amplitude modulation (PAM) chlorophyll fluorescence under atmospheric air or 100% N_2_ gas. In *T. japonicum* leaves, N_2_ gas treatment decreased the effective quantum yield of photosystem II (*Φ*_PSII_), which reflects the efficiency of photosynthetic electron transport (Maxwell and Johnson, 2000), at the end of the light period (phase IV) but not in the early light period (phase III). We assumed that in *T. japonicum* leaves at phase IV, when Rubisco mainly used atmospheric CO_2_, CO_2_ elimination from the air decreased the photosynthetic electron transport rate because of the deprivation of CO_2_ from the Calvin–Benson–Bassham reductive pentose phosphate cycle, the main sink of ATP and NADPH. By contrast, in the leaves at phase III, when Rubisco mainly used CO_2_ released from malate, CO_2_ elimination from the air did not cause lack of CO_2_ in photosynthetic cells and unaffected the *Φ*_PSII_ levels. Similar results were obtained in *T. aphyllum* roots but not in *T. japonicum* roots, where N_2_ gas treatment decreased *Φ*_PSII_ regardless of the light periods. The results of the PAM analysis were consistent with those of ^14^CO_2_ labeling analysis and malate determination, showing the availability of this CO_2_-elimination PAM (CE-PAM) analysis for characterizing the photosynthetic mode of CAM plants.

In this study, we performed CE-PAM analysis in well-known constitutive CAM plants, *Kalanchoe pinnata* and *Kalanchoe daigremontiana*. Moreover, we characterized four epiphytic orchids native to Japan (*Gastrochilus japonicus*, *Oberonia japonica*, *Bulbophyllum inconspicuum*, and *Bulbophyllum drymoglossum*), whose CO_2_ fixation pathways were previously unknown, along with *T. japonicum* and the C_3_ orchid *Cymbidium goeringii*. The CE-PAM analysis revealed that *G. japonicus*, *O. japonica*, and *B. inconspicuum* but not *B. drymoglossum* are constitutive CAM plants. To characterize *B. drymoglossum* in more detail, photosynthetic responses to water deficit or high light and leaf anatomy were investigated, revealing the peculiarity of *B. drymoglossum* in comparison with *B. inconspicuum*.

## Material & Methods

### Plant materials and growth conditions

All plants used in this study were grown at 23 °C under a 10 h light period (100□μmol photons m^−2^ s^−1^) between 9:00 and 19:00 in growth chambers, watered twice a week, unless otherwise stated.

*K. pinnata* and *K. daigremontiana* were provided by Dr. Kensuke Miyamoto, Professor emeritus of Osaka Prefecture University, and were grown to a height of ∼15 cm and ∼30 cm, respectively, on commercially available soil. For the CE-PAM analysis and titratable acidity assay, fully developed leaves (∼7 cm in length) of *K. pinnata* and those (∼8 cm in length) of *K. daigremontiana* were used as mature leaves. For both plants, the youngest or second-youngest leaves emerged from the apical shoot meristems, which were ∼2.5 cm in length, were used as immature leaves. *T. japonicum* plants were collected from a *Cryptomeria japonica*-dominated conifer plantation in Asakita, Hiroshima City, Hiroshima Prefecture, Japan, in November 2022. *G. japonicus* was collected from Kirishima City, Kagoshima Prefecture, Japan, in February 2021. *O. japonica* plants were collected from Gotenba City, Shizuoka Prefecture, Japan, in February 2021. *B. inconspicuum* and *B. drymoglossum* plants were collected from Miyakonojo City, Miyazaki Prefecture, Japan, in February 2021. *T. japonicum*, *O. japonica*, *B. inconspicuum*, and *B. drymoglossum* were grown on wet sphagnum moss, while *G. japonicus* was grown on naturally weathered, hollowed-out decayed wood. *C. goeringii* plants were collected from Hitachinaka City, Ibaraki Prefecture, Japan, in February 2021 and grown on commercially available acidic, well-drained soil.

To determine *Φ*_PSII_ and malate content in *B. drymoglossum* leaves under dry conditions, *B. drymoglossum* plants that had been grown on wet sphagnum moss were transferred onto dry sphagnum moss and grown for 2 days without watering. For malate determination in Fig. 7, leaves were sampled at 9:00 and 19:00 on the second day of the dry condition.

To determine the *Φ*_PSII_ levels of *B. drymoglossum* leaves under high light conditions, *B. drymoglossum* plants that had been grown under the 10-h period of 100□μmol photons m^−2^ s^−1^ light were transferred to the 10-h period of 600□μmol photons m^−2^ s^−1^ light and grown for 2 days.

### CE-PAM analysis

CE-PAM analysis was carried out using an IMAGING-PAM MAXI fluorometer (Walz, Effeltrich, Germany) and associated software (ImagingWin; Walz). Whole plants or leaves were enclosed in a gas chamber made from an airtight lunch box and a transparent lid. To determine stationary (*F*) and maximal chlorophyll fluorescence under light (*F*′_m_), plant samples in the gas chamber were illuminated with ∼0.2□μmol photons m^−2^ s^−1^ measuring light and 100□μmol photons m^−2^ s^−1^ actinic light for 15 min, with saturating pulse irradiated every 20 s. During the first 5 min of the experiment, atmospheric air was supplied to the gas chamber by an air pump at a flow rate of ∼60 ml s^-1^. Then, CO_2_-free air was supplied to the gas chamber from a high-pressure gas cylinder containing CO_2_-deprived (1<ppm) atmospheric air (G3, Taiyo Nippon Sanso JFP Corporation, Kawasaki, Japan) at a similar flow rate for 5 min, followed by re-supply of atmospheric air for 5 min. Before flowing into the gas chamber, the air was bubbled with deionized water to moisten the air. For the data in Fig. 3, leaves of *K. pinnata* and *K. daigremontiana* were dark-incubated for 30 min before measurement and minimal chlorophyll fluorescence (*F*_o_) and maximal chlorophyll fluorescence (*F*_m_) were determined before actinic light illumination. From the fluorescence yields acquired, *Φ*_PSII_ and other photosynthetic parameters were computed (Maxwell and Johnson, 2000; Guadagno *et al*., 2010).

### Titratable acidity

Mature and immature leaves of *K. pinnata* and *K. daigremontiana* (0.1∼1.5 g) were collected at the beginning (9:00) and the end of the light period (19:00), chopped with a razor blade after being weighed, frozen in liquid nitrogen and stored at –80 °C until use. Five milliliters of 20% (v/v) ethanol were added to the chopped leaves in a test tube, boiled for 10 min, and transferred to a new test tube. After this process was repeated one more time, the extracts were combined and titrated with 10 mM NaOH to an endpoint of pH 6.5 (Winter and Smith, 2022).

### Determination of malate content

Leaves of *C. goeringii*, *T. japonicum*, *G. japonicus*, *O. japonica*, *B. inconspicuum*, and *B. drymoglossum* were weighed and collected in microtubes at 9:00 and 19:00, immediately frozen in liquid nitrogen, and stored at –80 °C until use. The leaves were pulverized in liquid nitrogen, homogenized in 800 µL ice-cold 1 M HClO_4_, incubated on ice for 20 min, and centrifuged at 21,900 × *g* for 10 min. The supernatants were transferred to new microtubes and neutralized using 285 µL of 6M KOH. The content of L-malate in the neutralized solution was determined using an L-malic acid assay kit (Megazyme, Bray, Ireland), which includes L-malate dehydrogenase and glutamate oxaloacetate transaminase to metabolize L-malate and reduce NAD^+^ to NADH. The formation of NADH, which is stoichiometric with the amount of L-malate, was measured by the increase in absorbance at 340 nm with a V-730 BIO spectrophotometer.

### Leaf anatomy

Leaves of well-watered *B. inconspicuum* and *B. drymoglossum* were cross-sectioned to ∼60 μm thickness with a DTK-1000 microslicer (DOSAKA EM, Kyoto, Japan) or to ∼100 μm thickness with an automatic plant microtome MT-3 (NK system, Osaka, Japan) and observed under a BX51 light microscope connected to a DP21 digital camera (Olympus, Tokyo, Japan). The 60 μm thin sections were stained with toluidine blue O (0.05%, w/v) in 0.1 M phosphate buffer (pH 6.8) to observe structures of leaf parenchyma cells, whereas the 100 μm sections were observed without staining to examine the distribution of chloroplasts. Leaf thickness was determined by measuring the length between the adaxial and abaxial surfaces of the transverse sections of leaves with the ImageJ variant software Fiji (Schindelin *et al*., 2012). To determine the numbers of stomata, a thin layer of clear nail polish was applied on the adaxial and abaxial surface of leaves and was air-dried completely at room temperature. The nail polish impression was peeled off with clear tape and was observed under the BX51 light microscope. To determine the water content of leaves, fresh leaves from well-watered *Bulbophyllum* plants were weighed on an AUW120D microbalance (Shimadzu, Kyoto, Japan) to determine fresh weight and then dried at 60 °C until no further weight loss was observed. The lowest weight was determined as dry weight and water content (%) was calculated as follows: 100 × (fresh weight – dry weight) / fresh weight. To determine succulence of leaves, 4.03 mm^2^ discs were punched out from leaves of well-watered *Bulbophyllum* plants and were weighed. Succulence was computed as follows: (water content / 100) × (fresh weight / area).

## Results

### Mature leaves but not immature leaves of *K. pinnata* and *K. daigremontiana* show CAM-type ***Φ***_PSII_ responses to CO_2_ elimination

Ontogenetic shifts from C_3_ to CAM during leaf maturation are often observed in constitutive CAM plants (Winter, 2019). It is reported that fully developed leaves of *K. pinnata* and *K. daigremontiana* perform CAM photosynthesis, while their immature leaves mainly perform C_3_ photosynthesis (Winter *et al*., 2008; Nishida, 1978; Nishida *et al*., 1981). To test whether *Φ*_PSII_ responses to CO_2_ elimination reflect the ontogenetic C_3_-CAM shift during leaf development, we performed CE-PAM analysis in addition to titratable acidity measurement in immature and mature leaves of *K. pinnata* and *K. daigremontiana*. Plants were illuminated from 9:00 (dawn) to 19:00 (dusk) with ∼100 μmol photons m^−2^ s^−1^ every day. The *Φ*_PSII_ kinetics were determined at 11:00 and 19:00, whereas the titratable acidity was measured at 9:00 and 19:00. In both *K. pinnata* and *K. daigremontiana*, immature leaves showed decreased *Φ*_PSII_ levels in response to CO_2_ elimination and their recovery with CO_2_ re-addition both at 11:00 and 19:00, with *K. daigremontiana* having a greater *Φ*_PSII_ decrease than *K. pinnata* (Fig. 1A, B). The data indicate that the immature leaves of these species require atmospheric CO_2_ to perform efficient photosynthetic electron transport regardless of the light period, consistent with the low titratable acidity in these leaves even at dawn (Fig. 2). By contrast, mature leaves of both *K. pinnata* and *K. daigremontiana* showed no remarkable decrease in *Φ*_PSII_ levels at 11:00 (Fig. 1C, D), suggesting that these mature leaves use internal CO_2_ sources to drive photosynthetic electron transport in this period. However, at 19:00, mature leaves of *K. pinnata* and *K. daigremontiana* showed strong *Φ*_PSII_ decreases in response to CO_2_ elimination, consistent with the low titratable acidity at dusk (Fig. 2).

**Fig. 1.**
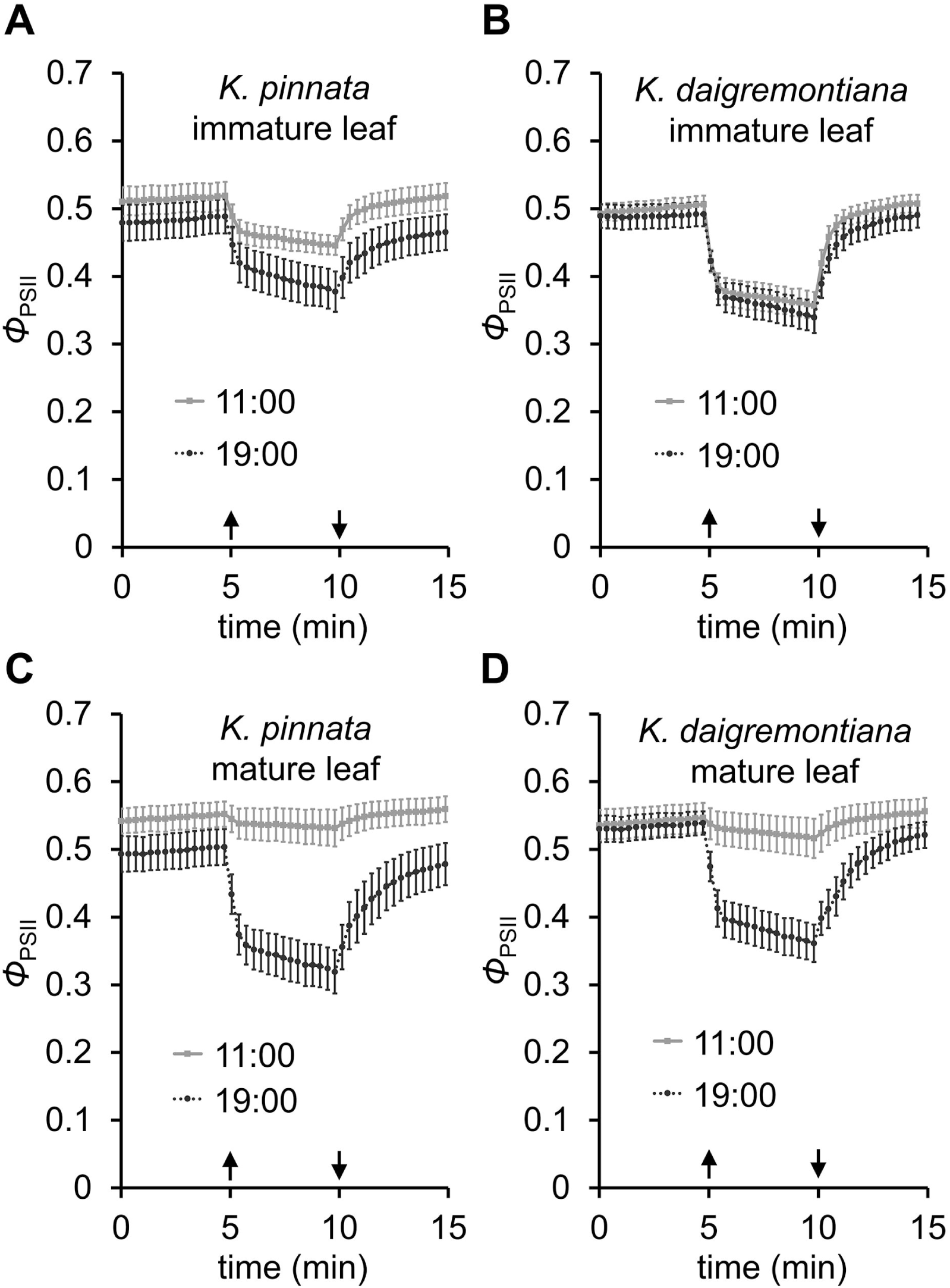
Fluctuation of *Φ*_PSII_ in response to CO_2_ elimination and re-addition in two *Kalanchoe* species. *Φ*_PSII_ levels at 2 h after the beginning of the light period (11:00) and the end of the light period (19:00) were determined under 100 μmol photons m^-2^ s^-1^ actinic light in immature (A, B) and mature leaves (C, D) of *Kalanchoe pinnata* (A, C) and *Kalanchoe daigremontiana* (B, D). Upward arrows indicate the switching of airflow from ambient air to CO_2_-eliminated air and downward arrows indicate the reversion of the flow from CO_2_-eliminated air to atmospheric air. Data are means ± SE from 9 leaves.

**Fig. 2.**
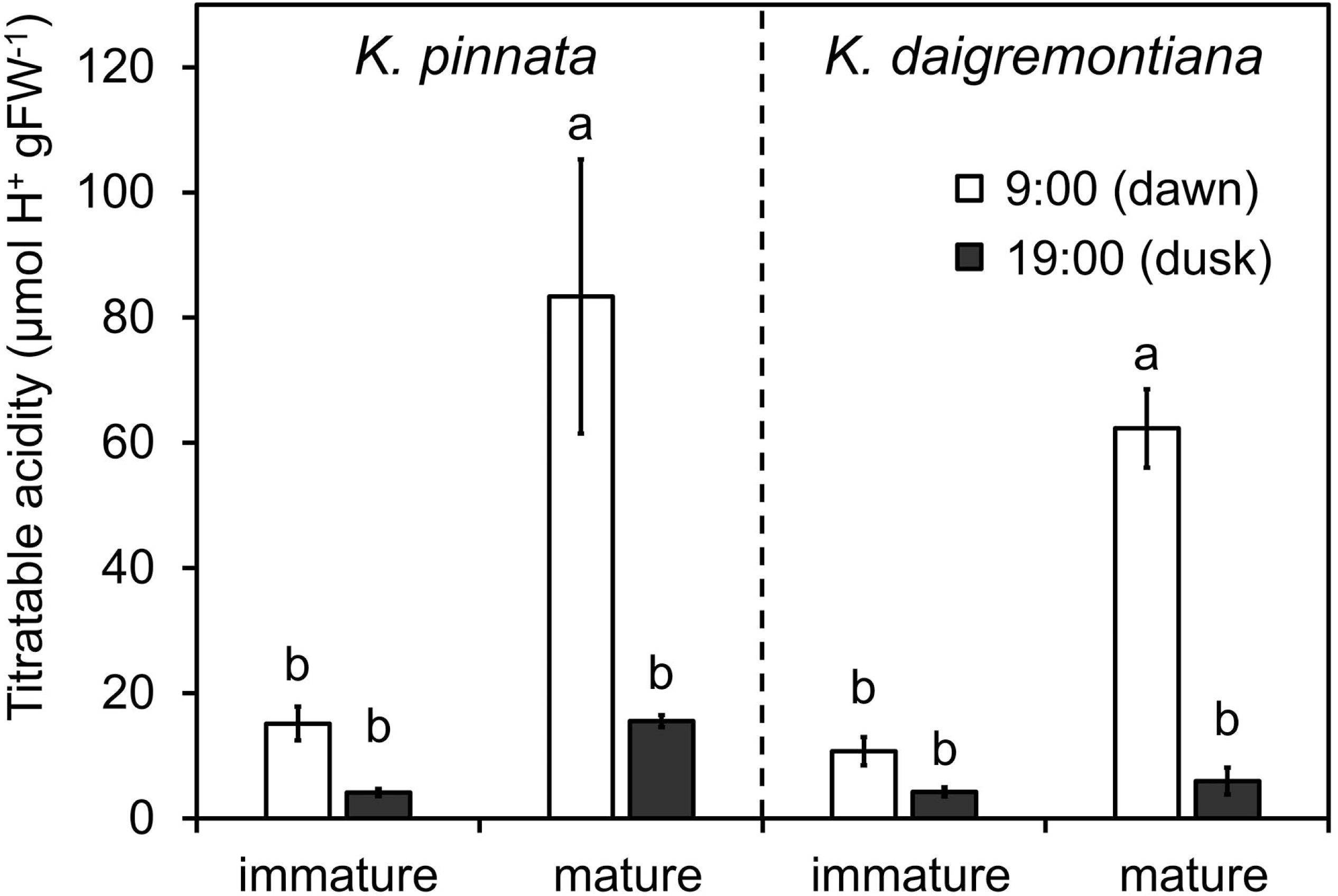
Titratable acidity of immature and mature leaves of *Kalanchoe pinnata* and *Kalanchoe daigremontiana* at dawn and dusk. Leaves were sampled at the beginning (9:00, dawn) and the end of the light period (19:00, dusk). Data are means ± SE from 3 independent leaves. The lowercase letters indicate statistically significant differences among all samples (P < 0.05, 1-way ANOVA and Tukey’s post hoc honestly significant difference test).

To explore the mechanism for the different *Φ*_PSII_ responses to CO_2_ elimination between immature and mature leaves of the *Kalanchoe* species, we analyzed the kinetics of several photosynthetic parameters (Maxwell and Johnson, 2000; Guadagno *et al*., 2010) in the CE-PAM analysis in immature and mature leaves of *K. daigremontiana* at 11:00 (Fig. 3). In immature leaves, *Φ*_NPQ_, which represents the yield of excess light energy dissipation in a regulated form mainly as heat, increased and decreased in response to CO_2_ elimination and re-addition, respectively. The fluctuation of *Φ*_NPQ_ was almost a mirror image of that of *Φ*_PSII_. By contrast, *Φ*_C_, which represents the yield of excess light energy dissipation in a non-regulated form mainly as fluorescence, only transiently increased and decreased at the onset of CO_2_ elimination and re-addition, respectively, in immature leaves. *Φ*_PSII_ can be viewed as a multiplication of the photochemical quenching coefficient (*q*P), which reflects the redox state of the plastoquinone pool and thus the openness of PSII, and *F′*_v_/*F′*_m_, which reflects the photochemical efficiency of the open PSII in an oxidized state. The *q*P level quickly decreased and increased in response to the elimination and re-addition of CO_2_, respectively, followed by a relatively slow decrease and increase of *F′*_v_/*F′*_m_. The nonphotochemical quenching (NPQ) parameter, *NPQ*, showed a fluctuation opposite to *F′*_v_/*F′*_m_. These data suggest that the CO_2_ elimination in immature *K. daigremontiana* leaves made PSII in a more reduced state, resulting in increased NPQ with decreased PSII photochemistry, which contributed to suppressing non-regulated dissipation of excess light energy (*Φ*_C_). In mature leaves, however, almost no change was observed in every photosynthetic parameter in response to CO_2_ elimination and re-addition, indicating that the photosynthetic electron transport functions independently of atmospheric CO_2_ in mature leaves of *K. daigremontiana* at phase III, consistent with its CAM activity.

**Fig. 3.**
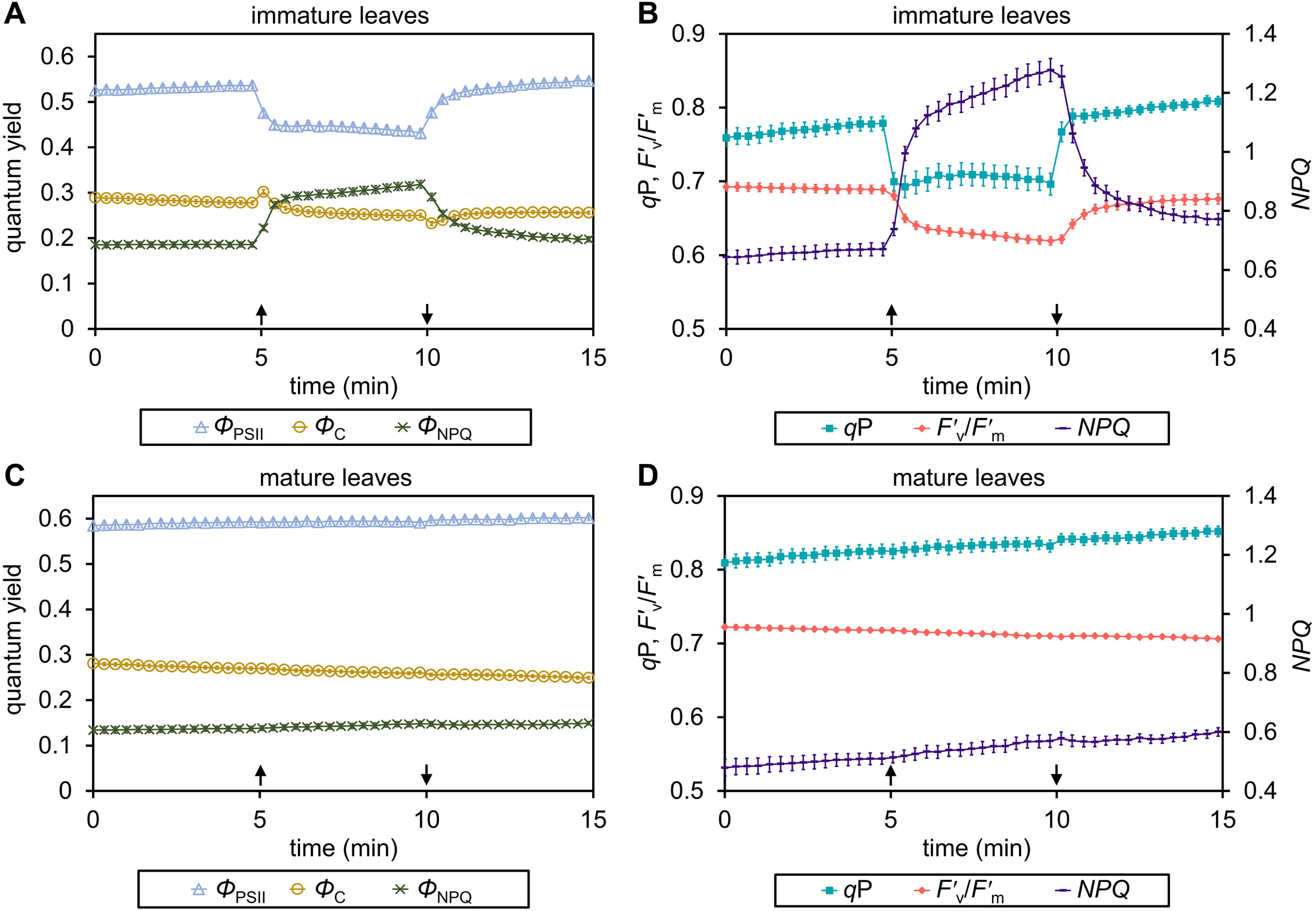
Changes in chlorophyll fluorescence parameters in response to CO_2_ availability in *Kalanchoe daigremontiana* leaves. Quantum yields of photosystem II (*Φ*_PSII_), nonregulated energy dissipation (*Φ*_C_), and regulated nonphotochemical quenching (*Φ*_NPQ_) (A, C) and the coefficient of photochemical quenching (*q*P), photochemical efficiency of open PSII (*F′*_v_/*F′*_m_), and the nonphotochemical quenching parameter (*NPQ*) (B, D) under 100 μmol photons m^-2^ s^-1^ actinic light were determined in immature leaves (A, B) and mature leaves (C, D) of *K. daigremontiana* at 2 h after the beginning of the light period (11:00). Upward arrows indicate switching of airflow from ambient air to CO_2_-eliminated air and downward arrows indicate the reversion of the flow from CO_2_-eliminated air to atmospheric air. Data are means ± SE from 12 leaves.

### Identification of CAM plants in epiphytic orchids by CE-PAM analysis

Our CE-PAM analysis in two *Kalanchoe* plants revealed CAM-associated *Φ*_PSII_ profiles, which are characterized by little change in phase III and a large reduction in phase IV in response to CO_2_ elimination. By using this CE-PAM technique, we assessed whether four epiphytic orchids native to Japan (*G. japonicus*, *O. japonica*, *B. inconspicuum*, and *B. drymoglossum*) perform CAM photosynthesis under well-watered conditions. Along with these plants, the terrestrial orchid *C. goeringii* was investigated as a typical C_3_ plant (Motomura *et al*., 2008), whereas *T. japonicum* was used as a control of CAM plants (Suetsugu *et al*., 2023). The *C. goeringii* leaves showed a strong decrease and increase in *Φ*_PSII_ levels in response to CO_2_ elimination and re-addition, respectively, regardless of the light period (Fig. 4), showing that *C. goeringii* always uses atmospheric CO_2_ for photosynthesis. By contrast, consistent with our previous study (Suetsugu *et al*., 2023), *T. japonicum* leaves showed the CAM-associated *Φ*_PSII_ kinetics; at 11:00, the *Φ*_PSII_ level was unresponsive to the changes in ambient CO_2_ concentrations, whereas *Φ*_PSII_ levels at 19:00 fluctuated in response to the atmospheric CO_2_ availability. The *Φ*_PSII_ kinetics of leaves of *G. japonicus*, *O. japonica*, and *B. inconspicuum* is similar to that of *T. japonicum* leaves, suggesting that these species constitutively perform CAM photosynthesis. Meanwhile, *B. drymoglossum*, a close relative of *B. inconspicuum*, showed strong *Φ*_PSII_ decreases with CO_2_ elimination both at 11:00 and 19:00, indicating that this plant continuously relies on atmospheric CO_2_ for efficient photosynthetic electron transport.

**Fig. 4.**
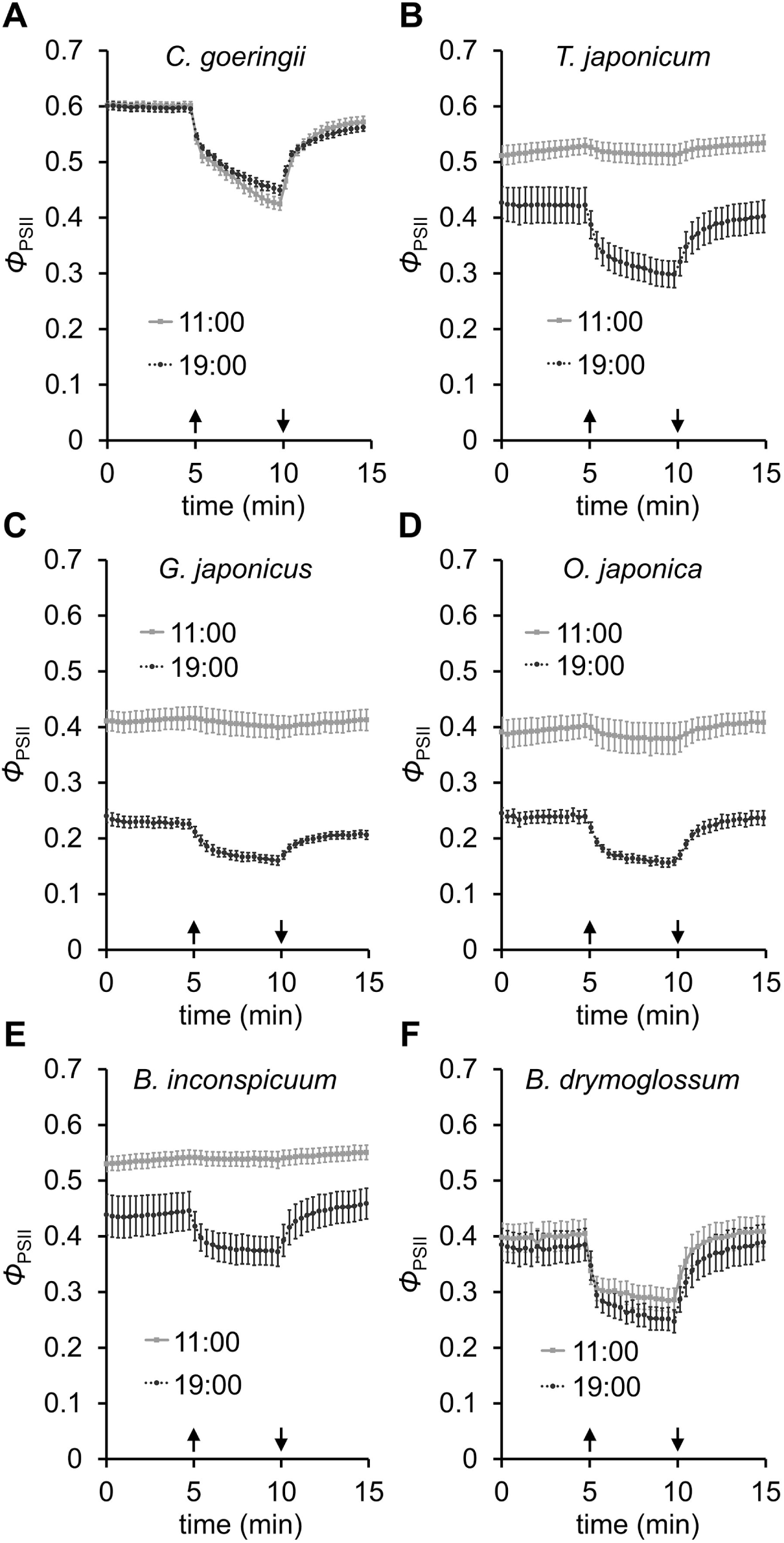
Responses of *Φ*_PSII_ to CO_2_ availability in orchids. *Φ*_PSII_ levels at 2 h after the beginning of the light period (11:00) and the end of the light period (19:00) were determined under 100 μmol photons m^-2^ s^-1^ actinic light in leaves of *Cymbidium goeringii* (A), *Thrixspermum japonicum* (B), *Gastrochilus japonicus* (C), *Oberonia japonica* (D), *Bulbophyllum inconspicuum* (E), and *Bulbophyllum drymoglossum* (F). Upward arrows indicate the switching of airflow from ambient air to CO_2_-eliminated air and downward arrows indicate the reversion of the flow from CO_2_-eliminated air to atmospheric air. Data are means ± SE from 5 leaves for each species.

To ascertain whether the *Φ*_PSII_ patterns of CE-PAM in these orchids are associated with the malate metabolism, we determined malate content at 9:00 (dawn) and 19:00 (dusk) (Fig. 5). Only a small amount of malate was detected in the leaves of *C. goeringii* both at 9:00 and 19:00, consistent with its C_3_ photosynthetic metabolism. By contrast, as previously observed (Suetsugu *et al*., 2023), malate content was high in *T. japonicum* leaves at 9:00 and the content was substantially decreased at 19:00, which is a typical feature of CAM plants. Similarly, the leaves of *G. japonicus*, *O. japonica*, and *B. inconspicuum* showed lower amounts of malate at 19:00 than at 9:00, so these plants consumed malate during the daytime as *T. japonicum* leaves did. Meanwhile, *B. drymoglossum* showed no significant decrease in malate content between 9:00 and 19:00. Our data demonstrate that *G. japonicus*, *O. japonica*, and *B. inconspicuum* but not *B. drymoglossum* are constitutive CAM plants, confirming the CE-PAM results.

**Fig. 5.**
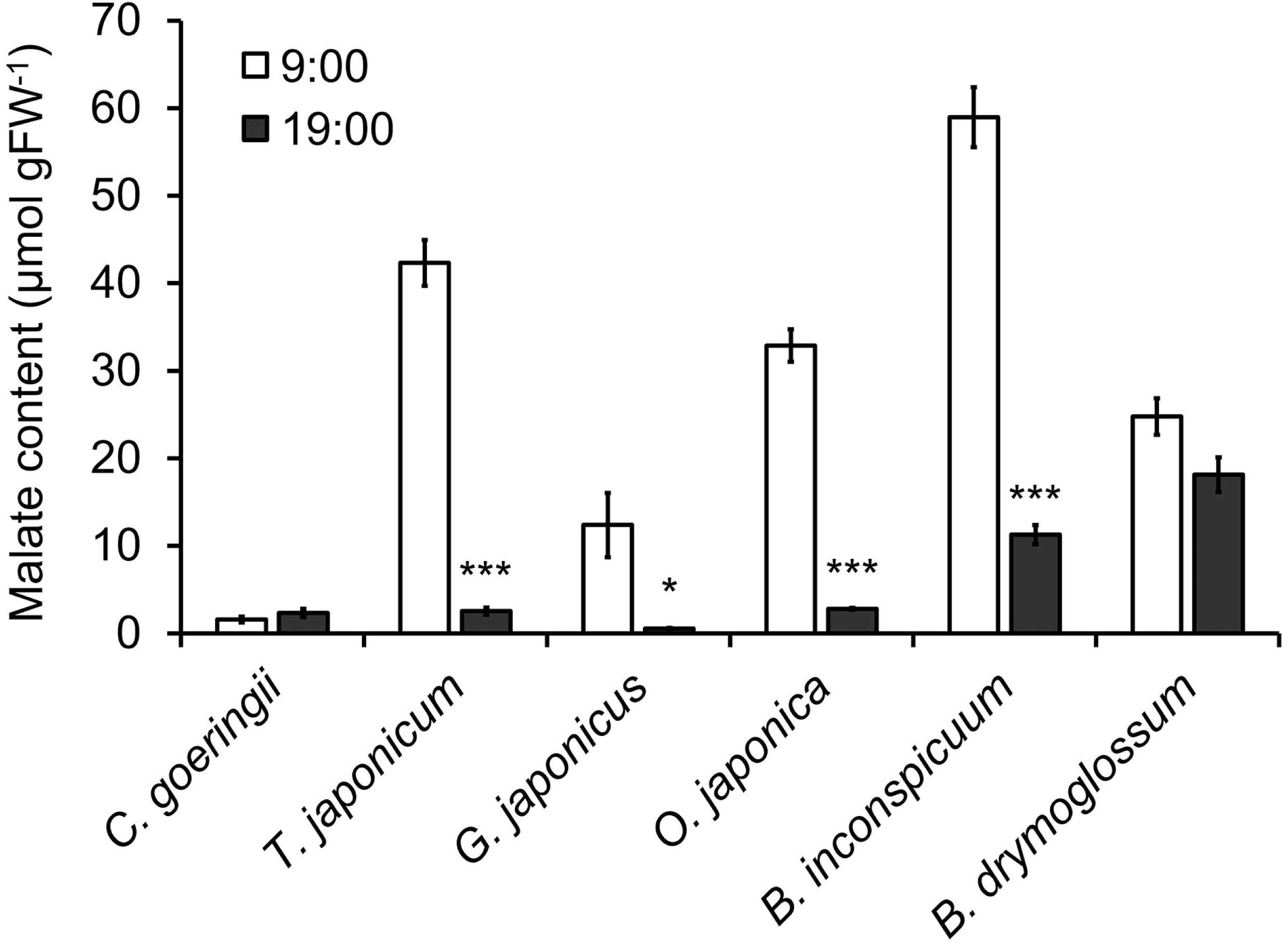
Malate content in orchids at dawn and dusk. Leaves of *Cymbidium goeringii*, *Thrixspermum japonicum*, *Gastrochilus japonicus*, *Oberonia japonica*, *Bulbophyllum inconspicuum*, and *Bulbophyllum drymoglossum* were sampled at the beginning (9:00) and the end of the light period (19:00). Data are means ± SE from 4 independent leaves. Asterisks show statistically significant differences between leaves at 9:00 and those at 19:00 for each species (P < 0.05, Welch’s t-test).

### Malate accumulated in *B. drymoglossum* leaves are not metabolized in response to drought and high light

Although *B. drymoglossum* did not show CAM-like characteristics in *Φ*_PSII_ patterns and malate metabolism under well-watered conditions (Figs. 4 and 5), this plant accumulated large amounts of malate at dawn and dusk. We hypothesized that *B. drymoglossum* leaves retain large amounts of malate under well-watered conditions to use it when they experience water deficit. To test this hypothesis, we performed CE-PAM analysis in *B. drymoglossum* plants deprived of water (Fig. 6). The *B. drymoglossum* plants grown on wet sphagnum moss were transferred onto dry sphagnum moss at 9:00 and grown for two days without watering. Before being transferred to the dry condition, the *Φ*_PSII_ of *B. drymoglossum* leaves strongly fluctuated in response to CO_2_ availability regardless of day and night, consistent with the data in Fig. 4. During the growth under the dry condition, the steady-state level of *Φ*_PSII_ in the atmospheric air decreased, presumably because of partial stomatal closure and a consequent decrease in CO_2_ uptake. Nevertheless, the water-deprived *B. drymoglossum* still showed *Φ*_PSII_ responses to CO_2_ availability at 11:00 similar to those at 19:00. The data imply that the water-deprived *B. drymoglossum* does not change their carbon metabolism from C_3_ to CAM in our experimental conditions.

**Fig. 6.**
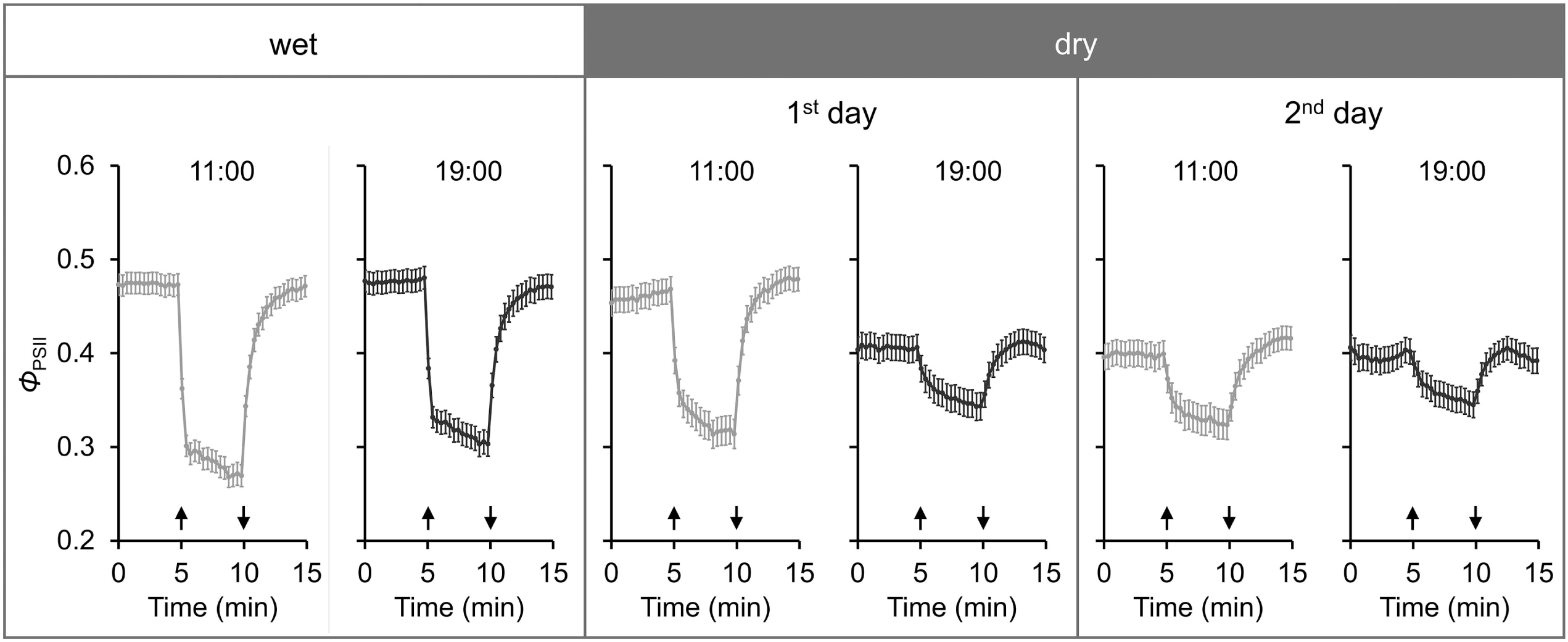
Effects of water deficit on *Φ*_PSII_ responses to CO_2_ availability in *Bulbophyllum drymoglossum*. *Φ*_PSII_ responses to CO_2_ elimination and re-addition under 100 μmol photons m^-2^ s^-1^ actinic light were determined at 2 h after the beginning of the light period (11:00) and the end of the light period (19:00). For wet condition control, *Φ*_PSII_ was determined in plants on wet sphagnum moss at 11:00 and 19:00. The plants were transferred onto dry sphagnum at 9:00 of the next day and grown for 2 days without watering, with *Φ*_PSII_ being determined at 11:00 and 19:00 each day. Data are means ± SE from 23 leaves. Upward arrows indicate the switching of airflow from ambient air to CO_2_-eliminated air and downward arrows indicate the reversion of the flow from CO_2_-eliminated air to atmospheric air.

To support the results of CE-PAM analysis, we placed *B. drymoglossum* plants on dry sphagnum moss and determined malate content in their leaves at 9:00 and 19:00 of the second day of water deprivation (Fig. 7). The result showed that the dry condition did not decrease the malate content in *B. drymoglossum* leaves compared with the wet condition, confirming that the malate stored in *B. drymoglossum* leaves is not used to cope with the sudden water deficit.

**Fig. 7.**
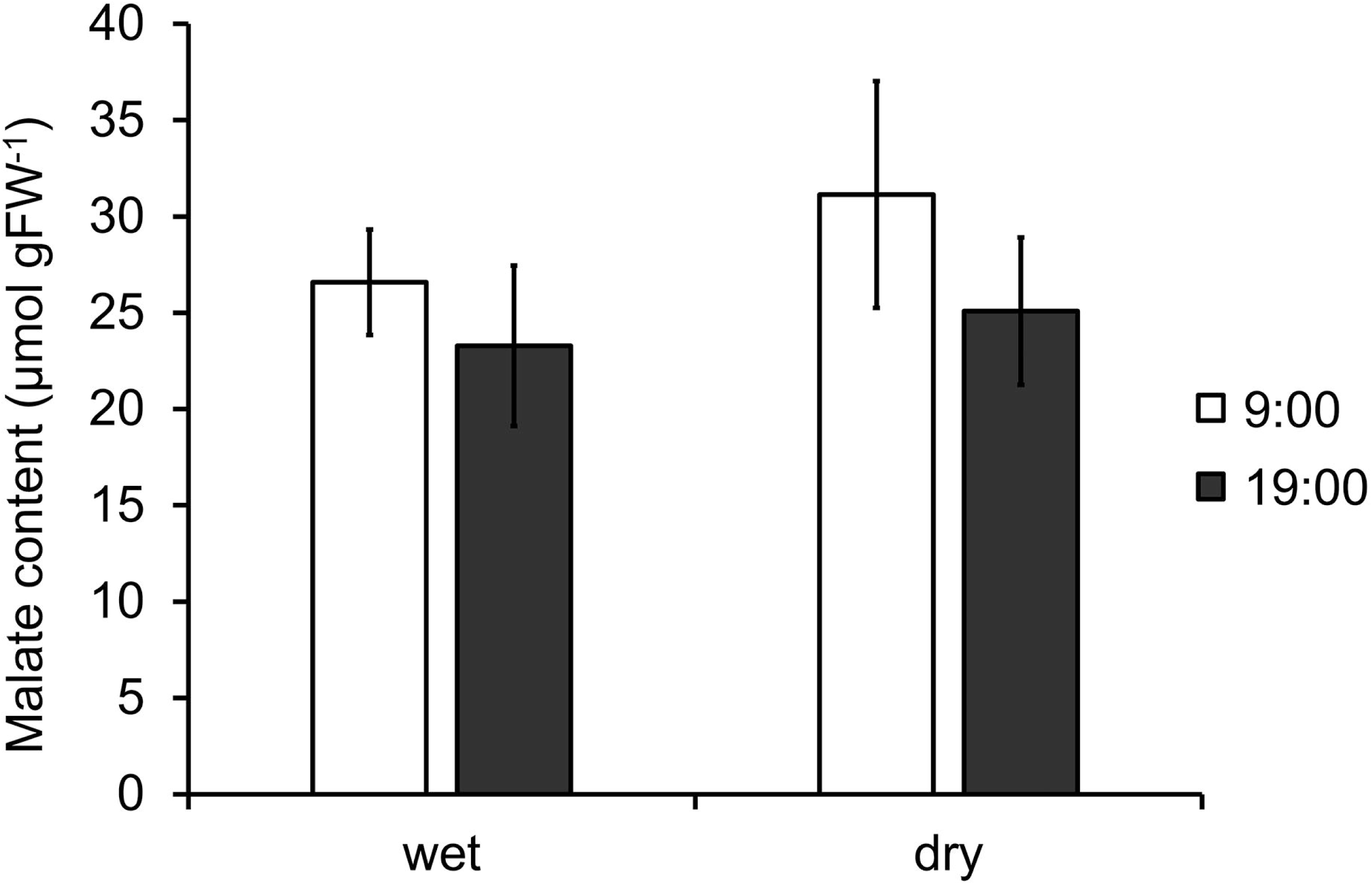
Stable malate content in leaves of *Bulbophyllum drymoglossum* under wet and dry conditions. Leaves of *B. drymoglossum* grown under wet or dry conditions were sampled at the beginning (9:00) and the end of the light period (19:00). Data are means ± SE from 3 independent leaves. No statistically significant differences were observed between leaves at 9:00 and those at 19:00 for each condition (P > 0.05, Welch’s t-test).

It is reported that leaves of *Clusia hilariana* change their photosynthetic mode from C_3_ to CAM in response to high light conditions; under the low light condition (200 μmol photons m^−2^ s^−1^), malate content in *C. hilariana* leaves was constant during day and night, but under the high light condition (∼700 μmol photons m^−2^ s^−1^), it decreased towards dawn and increased towards dusk, the typical characteristics of CAM plants (Miszalski *et al*., 2013). To assess whether the light intensity affects the photosynthetic mode of *B. drymoglossum* leaves, we treated *B. drymoglossum* plants with high light (600 μmol photons m^−2^ s^−1^) and performed CE-PAM analysis (Fig. 8). Plants grown on wet sphagnum moss under 100 μmol photons m^−2^ s^−1^ light were exposed to 600 μmol photons m^−2^ s^−1^ light between 9:00 and 19:00 on the same mat for 2 days. Although the overall *Φ*_PSII_ levels were decreased by the high light treatment, presumably because of the development of NPQ, the response to CO_2_ was continuously observed. The data suggest that the high light irradiation did not change the carbon assimilation metabolism in *B. drymoglossum*.

**Fig. 8.**
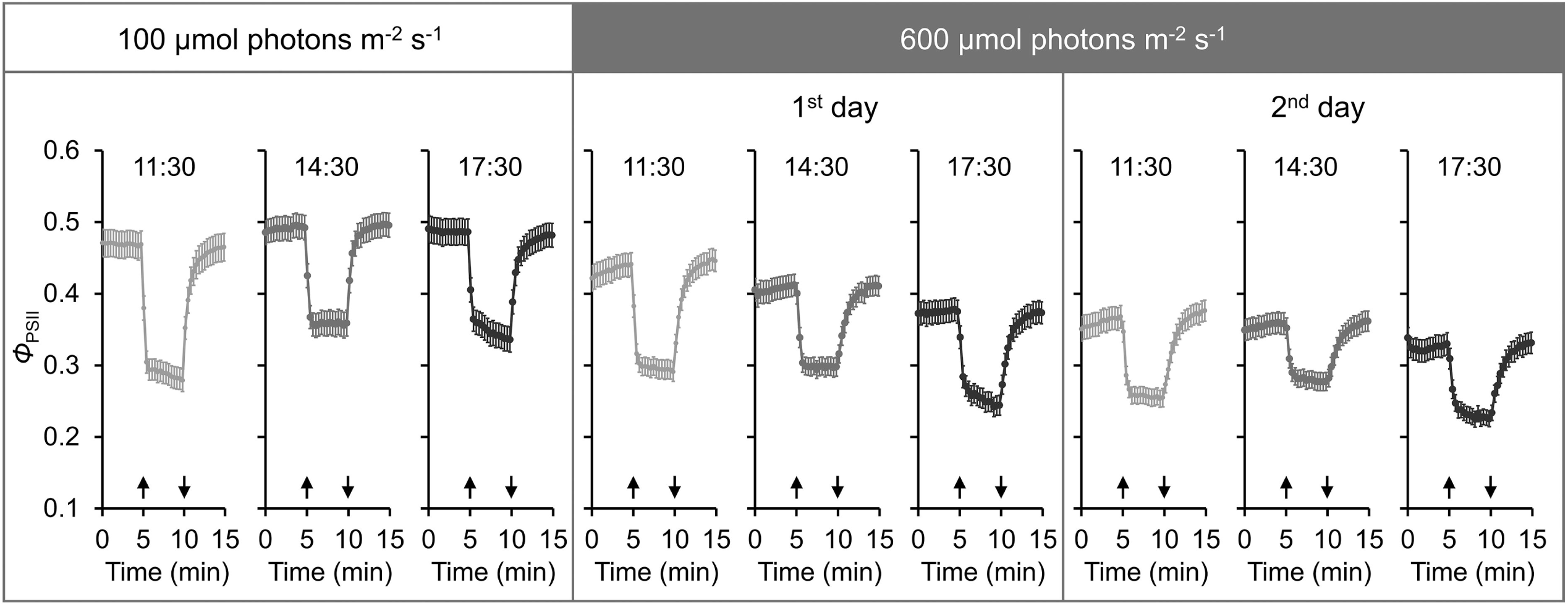
Effects of high light on *Φ*_PSII_ responses to CO_2_ availability in *Bulbophyllum drymoglossum*. *Φ*_PSII_ responses to CO_2_ elimination and re-addition under 100 μmol photons m^-2^ s^-1^ actinic light were determined at 11:30, 14:30, and 17:30. For low light control, *Φ*_PSII_ was determined in plants that had been grown under the 10-h period of 100 μmol photons m^-2^ s^-1^ white light. Then the plants were grown under the 10-h period of 600 photons m^-2^ s^-1^ white light and *Φ*_PSII_ was determined at the indicated time during the growth. Data are means ± SE from 20 leaves. Upward arrows indicate the switching of airflow from ambient air to CO_2_-eliminated air and downward arrows indicate the reversion of the flow from CO_2_-eliminated air to atmospheric air.

### Anatomical characterization of *B. drymoglossum* leaves compared to *B. inconspicuum* leaves

Photosynthetic organs of CAM species are known to share anatomical traits that reflect functional constraints associated with their specialized CO_2_ metabolism, with thick and succulent leaves being one of the general traits widely observed in CAM species (Gilman *et al*., 2024). To assess whether the leaf anatomy of non-CAM *B. drymoglossum* differs from that of CAM *B. inconspicuum*, we compared leaf structures between these two *Bulbophyllum* species (Table 1, Fig. 9).

**Fig. 9.**
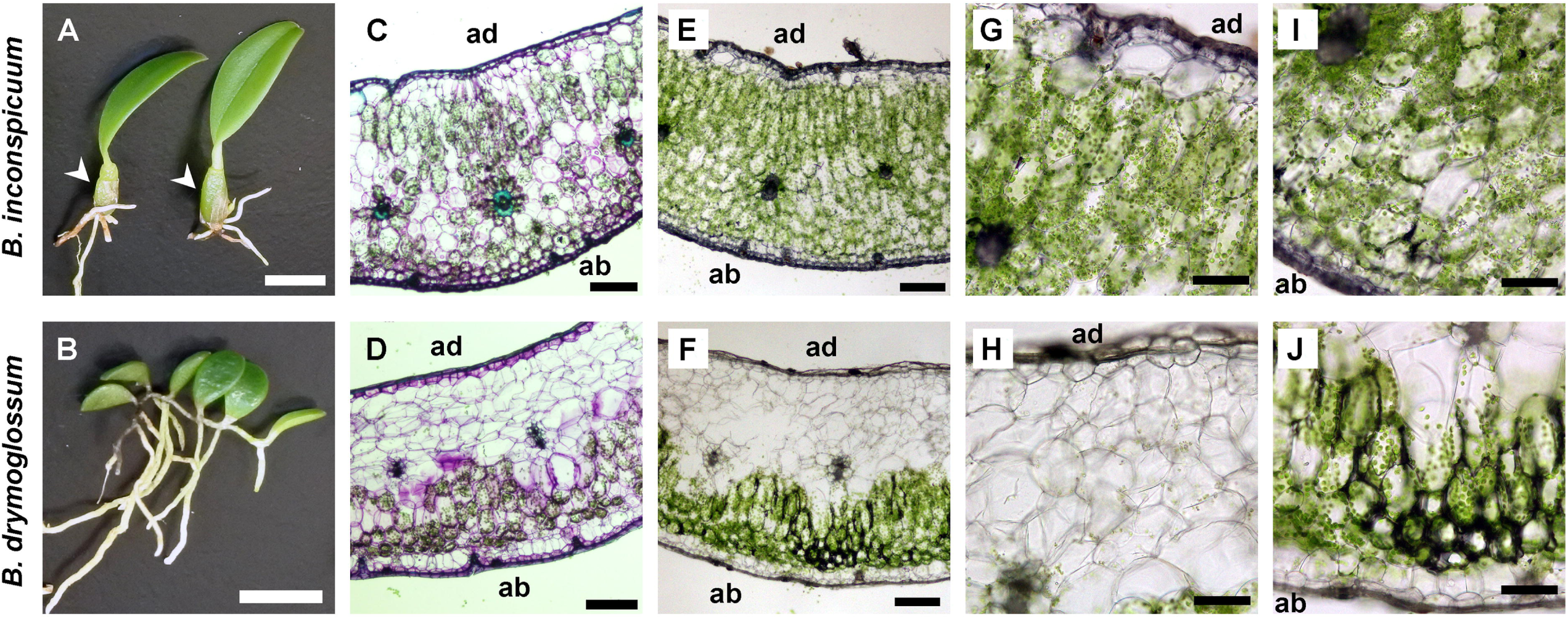
Leaf anatomy of *Bulbophyllum inconspicuum and Bulbophyllum drymoglossum*. (A, B) Specimens of *B. inconspicuum* (A) and *B. drymoglossum* (B) excised from each colony grown on sphagnum moss. Arrowheads indicate pseudobulbs developed with leaves in *B. inconspicuum*. No such structures were observed in *B. drymoglossum*. Bars = 5 mm. (C-F) Toluidine blue-stained (C, D) and unstained (E, F) transverse sections of leaves from *B. inconspicuum* (C, E) and *B. drymoglossum* (D, F). Bars = 200 μm. ad and ab, adaxial and abaxial side of leaves, respectively. (G-J) Close-up images of the ad (G, H) and ab (I, J) sides of leaves from *B. inconspicuum* (G, I) and *B. drymoglossum* (H, J). Bars = 100 μm.

**Table 1.** Anatomical features of leaves of Bulbophyllum inconspicuum and Bulbophyllum drymoglossum.

| | Thickness<br>($\mu\text{m}$) | Number of stomata ($\text{mm}^{-2}$) | | Water content<br>(%) | Succulence<br>($\text{kg m}^{-2}$) |
| --- | --- | --- | --- | --- | --- |
|  |  | Abaxial<br>surface | Adaxial<br>surface |  |  |
| *B. inconspicuum* | $882 \pm 37$ (n = 4) | $58.5 \pm 4.4$ (n = 3) | Not detected | $87.4 \pm 0.4$ (n = 9) | $0.745 \pm 0.016$ (n = 5) |
| *B. drymoglossum* | $792 \pm 17$ (n = 4) | $35.1 \pm 4.7$ (n = 3) | Not detected | $92.2 \pm 0.7$ (n = 9) | $0.804 \pm 0.028$ (n = 5) |

Data are mean ± SE.

Observation of transverse sections of fully developed leaves revealed that *B. inconspicuum* leaves (882 ± 37 μm) were slightly thicker than *B. drymoglossum* leaves (792 ± 17 μm). In the adaxial side of the *B. inconspicuum* leaves, vertically elongated mesophyll cells containing many chloroplasts formed 2-3 cell layers below a vacuolated achlorophyllous cell layer, which would be water storage parenchyma called hydrenchyma (Males, 2017) (Fig. 9). In the abaxial side of the *B. drymoglossum* leaves, spherical mesophyll cells with relatively fewer chloroplasts were mainly observed, with large mesophyll cells observed above the abaxial spherical mesophyll cells. The palisade-like layers on the adaxial side and spongy-like layers on the abaxial side together formed the large chlorenchyma in *B. inconspicuum* leaves. By contrast, in *B. drymoglossum* leaves, large vacuolated cells lacking chloroplasts occupied more than half of the transversal area from the adaxial side, forming large achlorophyllous hydrenchyma layers. In the abaxial side of the *B. drymoglossum* leaves, 1-2 cell layers of vertically elongated mesophyll cells containing many chloroplasts were observed above the spherical mesophyll cells. The number of stomata on the abaxial surface (35.1 ± 4.7 mm^-2^) of *B. drymoglossum* leaves was fewer than that of *B. inconspicuum* (58.5 ± 4.4 mm^-2^), with no obvious stomatal structures being detected on the adaxial leaf surface of both species.

*B. inconspicuum* develops a pseudobulb, which is known to serve as a water storage organ, at the bottom of a leaf, while *B. drymoglossum* has no such obvious water storage organs with its leaves (Fig. 8). We hypothesized that the large achlorophyllous hydrenchyma tissue occupying the upper half of the *B. drymoglossum* leaves, which is composed of large vacuolated cells, may increase the water storage capacity in this species. To support this hypothesis, we compared the water content in the leaves of *B. drymoglossum* with that of *B. inconspicuum*. The percentage of water content in fresh leaves of well-watered plants was 92.2 ± 0.7 % for *B. drymoglossum*, which was higher than 87.4 ± 0.4 % for *B. inconspicuum*. We also determined leaf succulence (water content per leaf area) in these two *Bulbophyllum* species. The leaf succulence of *B. drymoglossum* (0.804 ± 0.028 kg m^-2^) was slightly higher than that of *B. inconspicuum* (0.745 ± 0.016 kg m^-2^). These results indicate that *B. drymoglossum* leaves have a higher water storage capacity than *B. inconspicuum* leaves.

## Discussion

CAM plants are characterized by their ability to accumulate malate through fixation of atmospheric CO_2_ during the night (phase I) and to decarboxylate it to provide CO_2_ to Rubisco during the daytime (phase III) (Osmond, 1978). Accordingly, in phase III, they do not rely on atmospheric CO_2_ to drive the reductive pentose phosphate cycle as a sink for ATP and NADPH produced via photosynthetic electron transport. However, in phase IV, when the malate stored in vacuoles is depleted, CAM plants open stomata and use atmospheric CO_2_ for the reductive pentose phosphate cycle. These metabolic characteristics were reflected by different *Φ*_PSII_ responses to atmospheric CO_2_ availability between phase III and phase IV. In phase III, *Φ*_PSII_ in mature leaves of *K. pinnata* and *K. daigremontiana* was unaffected by CO_2_ concentration in the ambient air, whereas in phase IV, it quickly decreased and increased in response to CO_2_ elimination and re-addition, respectively (Fig. 1C, D). Meanwhile, *Φ*_PSII_ in young leaves of these two *Kalanchoe* species, which are known to perform C_3_ photosynthesis (Winter *et al*., 2008; Nishida, 1978; Nishida *et al*., 1981), is fluctuated in response to atmospheric CO_2_ availability regardless of light periods (Fig. 1A, B). Therefore, the responsiveness of *Φ*_PSII_ to CO_2_ availability at daytime can be used to distinguish C_3_ and CAM photosynthesis, and in fact, we identified *G. japonicus*, *O. japonica*, and *B. inconspicuum* as constitutive CAM plants and *B. drymoglossum* as a C_3_ plant (Fig. 4 and 5). Our CE-PAM analysis provides a novel approach to identify and characterize CAM plants easily and quickly with simple experimental procedures. Imaging-based PAM chlorophyll fluorescence analysis would enable large-scale screening of plants and mutants performing CAM or C_3_ photosynthesis under various growth conditions.

The changes in *Φ*_PSII_ in response to atmospheric CO_2_ availability during C_3_ photosynthesis were accompanied by changes in several other photosynthetic parameters (Fig. 3A, B). *q*P showed acute responses to CO_2_ concentration in immature *K. daigremontiana* leaves, suggesting that, in C_3_-performing plants, CO_2_ elimination from the ambient air quickly reduces the plastoquinone pool and CO_2_ re-addition reversely oxidizes it. Following the acute changes in *q*P, NPQ is developed by CO_2_ elimination and relaxed by CO_2_ re-addition. The rapid development and relaxation of the *NPQ* parameter within 1 min indicate that the energy-dependent quenching (*q*E), which is the most rapidly inducible NPQ component triggered by the ΔpH generation across the thylakoid membrane (Murchie and Ruban, 2020), is the main component of the NPQ mechanism developed by CO_2_ elimination. Interruption of the reductive pentose phosphate cycle and consequent attenuation of ATP consumption by CO_2_ elimination would cause acidification of the thylakoid lumen and thereby induce the *q*E-dependent NPQ (Kuvykin *et al*., 2011). Considering that the *Φ*_C_, which reflects the non-regulated energy dissipation (Guadagno *et al*., 2010), is only slightly and transiently increased with CO_2_ elimination, the activation of NPQ would contribute to minimizing harmful energy dissipation caused by retarded photosynthetic electron transport with CO_2_ limitation. Of note, certain levels of *Φ*_PSII_ are maintained during the CO_2_ elimination in leaves performing C_3_ photosynthesis, including leaves of *C. goeringii* and *B. drymoglossum*, immature leaves of *K. daigremontiana*, and leaves of CAM plants at phase IV. Because we filled leaves with 79% N_2_ and 21% O_2_ gas to eliminate CO_2_, O_2_ in the air may function as a sink of electrons from photosynthetic electron transport in addition to other electron-accepting metabolisms in chloroplasts (Kuvykin *et al*., 2011).

In contrast to the rapid and strong decrease in *Φ*_PSII_ in response to CO_2_ elimination in leaves performing C_3_ photosynthesis, leaves performing CAM photosynthesis maintained high *Φ*_PSII_ levels under the CO_2_-eliminated air condition. In mature *K. daigremontiana* leaves in phase III, no rapid *NPQ* development was observed, with the *q*P levels being constant (Fig. 3C, D), demonstrating that the supply of CO_2_ from malate functions to maintain high photosynthetic electron transport activity with suppressing NPQ mechanisms. The elimination of CO_2_ from the ambient air mimics the conditions that force leaves to close their stomata such as severe drying conditions. Accumulating malate allows leaves to maintain the plastoquinone pool in an oxidized state even with their stomata being closed, which contributes to increasing photosynthetic production while avoiding photoinhibition. Adams III and Osmond (1988) reported that *K. pinnata* leaves experienced severe photoinhibition with high light in the absence of CO_2_ when malate accumulation was prevented during the previous dark period. These data indicate that the malate accumulated in leaf cells contributes to preventing photoinhibition when the CO_2_ uptake from the atmosphere is limited under high light. Nevertheless, in CAM plants, the high photosynthetic electron transport activity behind closed stomata at phase III strongly increases internal O_2_ concentrations and causes the formation of reactive oxygen species, which may be a reason why CAM plants have developed effective antioxidative response systems (Lüttge, 2004).

We revealed that *B. drymoglossum* contained a high amount of malate in leaves regardless of the light period (Fig. 5), which led us to hypothesize that this species uses malate when it experiences drought stress. However, our CE-PAM analysis and malate determination indicated that *B. drymoglossum* leaves did not change their photosynthetic mode to CAM at least within 2 d of water deprivation (Fig. 6, 7). We also tested high light as a possible inducer of CAM photosynthesis in *B. drymoglossum*. However, the high light of 600 μmol photons m^−2^ s^−1^ did not make this plant change its C_3_-type *Φ*_PSII_ responses (Fig. 8). Currently, we do not know the reason why *B. drymoglossum* constantly accumulates malate in its leaves. Mobilization of malate requires the transport of malate from vacuoles and its decarboxylation by specific enzymes including NAD(P)-malic enzyme and phosphoenolpyruvate carboxykinase (Ceusters *et al*., 2021). Further investigations will be required to address the mechanisms that determine the availability of malate in *B. drymoglossum* leaves. Of note, *B. inconspicuum*, a close relative of *B. drymoglossum*, had relatively high malate content at dusk compared with other epiphytic CAM orchids investigated in this study (Fig. 5), although *B. inconspicuum* showed C_3_-like *Φ*_PSII_ fluctuation in response to CO_2_ elimination at dusk (Fig. 4). These data indicate that *B. inconspicuum* leaves switch their photosynthetic mode from CAM to C_3_ while retaining a relatively high amount of malate. It was reported in *Phalaenopsis* “Sacramento” that the length of phase III and the start of phase IV were adjusted to the moment when malate was depleted, suggesting that the cellular malate content is linked with the transition from phase III to phase IV (Hogewoning *et al*., 2021). However, under low light conditions, the *Phalaenopsis* plants shifted to phase IV even though they retained a high amount of malate. Thus, the transition from phase III to phase IV is regulated in a complex manner, which has not yet been well understood (Ceusters *et al*., 2021). Some *Clusia* species expressing CAM are reported to show high titratable acidity even at dusk (Winter and Smith, 2022). Moreover, even non-CAM *Clusia* species, which showed no diel fluctuations of titratable acidity, had high values of tissue acidity at dawn and dusk. Therefore, the capacity to accumulate and mobilize malate would vary depending on plant species. Similar to *Clusia* species, *Bulbophyllum* species may generally have a high capacity to accumulate malate that is not used for CAM activity.

Thick, succulent photosynthetic tissues are frequently observed in CAM plants (Herrera, 2020; Gilman *et al*., 2024). Although the leaves of *B. drymoglossum* were slightly thinner than *B. inconspicuum* (Table 1), they were thicker than those of typical C_3_ plants with similar leaf sizes such as *Arabidopsis thaliana* (150 ∼ 200 μm of leaf thickness) (Wuyts *et al*., 2012). The succulence and the leaf thickness indicate that leaves of both *B. drymoglossum* and *B. inconspicuum* have a characteristic similar to succulent CAM plants (Barrera Zambrano *et al*., 2014; Heyduk *et al*., 2016; Gilman *et al*., 2024). Nevertheless, unlike *B. inconspicuum*, *B. drymoglossum* showed no CAM features in malate metabolism and photosynthetic CO_2_ responses. Except for some CAM groups, leaf thickness and other anatomical features of leaves are not strongly correlated with the degree of CAM expression (Herrera, 2020). Although the leaf thickness was relatively similar between *B. inconspicuum* and *B. drymoglossum*, their leaf anatomy was largely different; *B. drymoglossum* leaves have large chloroplast-lacking cell layers, namely the achlorophyllous hydrenchyma tissue, in the adaxial half, whereas *B. inconspicuum* leaves develop chloroplasts across the adaxial and abaxial sides, with only single hydrenchyma cell layer observed under the adaxial surface (Fig. 9). Considering that *B. drymoglossum* leaves have no external water storage organs in contrast to *B. inconspicuum* leaves that are connected to pseudobulbs, the large achlorophyllous hydrenchyma tissue in *B. drymoglossum* leaves would serve as the main site for water storage. Of note, achlorophyllous hydrenchyma tissues are observed in a wide range of succulent plants (Males, 2017), and no or little correlation has been indicated between hydrenchyma thickness and CAM expression (Barrera Zambrano *et al*., 2014; Martin *et al*., 2019; Males, 2018), consistent with our observations in *B. drymoglossum*. The physiological significance of the development of the large achlorophyllous parenchyma tissue and its evolutionary relationships with other characteristics such as the lack of pseudobulbs and non-CAM photosynthesis in *B. drymoglossum* leaves need to be elucidated in future studies.

## Author contributions

SB performed experiments and analyzed the data; KS collected plant materials and edited the manuscript; KK conceived the project, designed and performed experiments, analyzed the data, and wrote the manuscript. All the authors reviewed the manuscript and approved the final version of the manuscript.

## Data availability

Data will be made available from the corresponding author upon request.

## Abbreviations

CAM: Crassulacean acid metabolism
CE-PAM: CO_2_-elimination PAM
*F′*_v_/*F′*_m_: photochemical efficiency of the open PSII in an oxidized state
NPQ: nonphotochemical quenching
PAM: pulse amplitude modulation
PEPC: phosphoenolpyruvate carboxylase
*q*P: photochemical quenching coefficient
*Φ*_C_: yield of excess light energy dissipation in a non-regulated form mainly as fluorescence
*Φ*_PSII_: effective quantum yield of photosystem II
*Φ*_NPQ_: yield of excess light energy dissipation in a regulated form mainly as heat

